# Rere-dependent Retinoic Acid signaling controls brain asymmetry and handedness

**DOI:** 10.1101/578625

**Authors:** Michael Rebagliati, Gonçalo C. Vilhais-Neto, Alexandra Petiet, Merlin Lange, Arthur Michaut, Jean-Luc Plassat, Julien Vermot, Fabrice Riet, Vincent Noblet, David Brasse, Patrice Laquerrière, Delphine Cussigh, Sébastien Bedu, Nicolas Dray, Mohamed Sayed Gomaa, Claire Simons, Hamid Meziane, Stéphane Lehéricy, Laure Bally-Cuif, Olivier Pourquié

## Abstract

While the vertebrate brain appears largely bilaterally symmetrical in humans, it presents local morphological Left-Right (LR) asymmetries as, for instance, in the *petalia*. Moreover, higher functions such as speech or handedness are asymmetrically localized in the cortex. How these brain asymmetries are generated remains unknown. Here, we reveal a striking parallel between the control of bilateral symmetry in the brain and in the precursors of vertebrae called somites, where a “default” asymmetry is buffered by Retinoic Acid (RA) signaling. This mechanism is evident in zebrafish and mouse and, when perturbed in both species, it translates in the brain into lateralized alterations of patterning, neuronal differentiation and behavior. We demonstrate that altering levels of the mouse RA coactivator Rere results in subtle cortex asymmetry and profoundly altered handedness, linking patterning and function in the motor cortex. Together our data uncover a novel mechanism that could underlie the establishment of brain asymmetries and handedness in vertebrates.

---

The bilateral symmetry of vertebrates is often assumed to be a default state of embryonic organization, but how developmental programs are coordinated to ensure that the left side develops as a mirror image of the right side is mostly unknown. The formation of somites provides an excellent illustration of the LR coordination of patterning processes in the vertebrate embryo. Somites form synchronously on the left and right side from the presomitic mesoderm (PSM) and they differentiate in a synchronized manner to produce the bilaterally symmetrical vertebral column. The LR symmetry of somite formation is controlled by a bilaterally symmetrical gradient of Fibroblast Growth Factor (FGF) signaling peaking in the posterior PSM ^1,2^. This FGF gradient is antagonized by RA signaling produced by the forming somites anteriorly ^3–4^. Strikingly, in RA-deficient mice lacking the RA biosynthetic enzyme Raldh2, the *Fgf8* gradient in the PSM becomes asymmetric, resulting in a delay of somitogenesis in the right side^5–6^. A similar phenotype is observed in mice mutant for the RA coactivator Rere, in which RA signaling is decreased ^1^. In the viscera, bilateral coordination of the development of the two embryonic sides is altered in response to a LR signaling pathway that controls their asymmetric development ^8^. In mouse and zebrafish, an Fgf8-dependent signal derived from the Node activates Nodal in the left lateral plate to specify the left embryonic identity^9–10^. Mutation of *Fgf8* or *Nodal* can result in *situs inversus* where visceral organs become organized in a mirror-image fashion ^8–10^. The right-sided somitogenesis defect observed in RA-deficient mice can be reversed by crossing them with the *inversus viscerum* (*iv*) mutant mice in which embryos exhibit *situs inversus* ^7,11^. Together, these experiments suggest that Rere-dependent RA signaling interacts with FGF to buffer the action of the LR pathway and maintain the bilateral symmetry of somites. The molecular mechanism underlying the interaction between RA and FGF signaling which maintains somites symmetry is currently unknown. Whether the role of RA in bilateral symmetry extends beyond somites has not been investigated.

Remarkably, the hindbrain and cortex are also patterned by sets of opposing bilateral gradients of FGF and RA signaling, raising the possibility that these pathways might also be implicated in the control of bilateral symmetry in these structures ^12,13^. The posterior gradient of RA involved in hindbrain patterning arises from the somites and it interacts with FGF signaling from the midbrain-hindbrain (isthmic) organizer to specify the identity of the forming rhombomeres ^14^. To explore whether RA is also involved in the control of bilateral symmetry of the hindbrain, we incubated zebrafish embryos beginning at the 8-128 cell stage (prior to 4h post-fertilization) with the pan-RA receptor antagonist, BMS-204493 (BMS)^15^. We first examined expression of the segmentation gene *deltaC*, which is expressed in a bilaterally symmetrical fashion in the PSM ^16^. As expected, BMS treatment resulted in expression of *deltaC* becoming asymmetric between the left and right PSM, suggesting that LR synchronization of somite segmentation is disrupted (Fig. 1a-b). We also observed striking bilateral symmetry defects in the hindbrain using the rhombomere (r) 3 and 5 marker *krox20* (*egr2a*) (Fig. 1c-e, Supplementary Table 1, Extended data figure 1a-f). In r5, asymmetries were variable in pattern but present as a left or right hemi-segmental absence of *kro×20*expression in the extreme (Fig. 1c-d, Extended data Fig. 1a-f). The *kro×20* r3 stripe was not affected. BMS treatment also reproduced the previously published *krox20*expression phenotypes resulting from hindbrain anteriorization, including a complete loss of r5 expression (Extended data Figure 1c-d)^17–20^.

**Figure 1:**
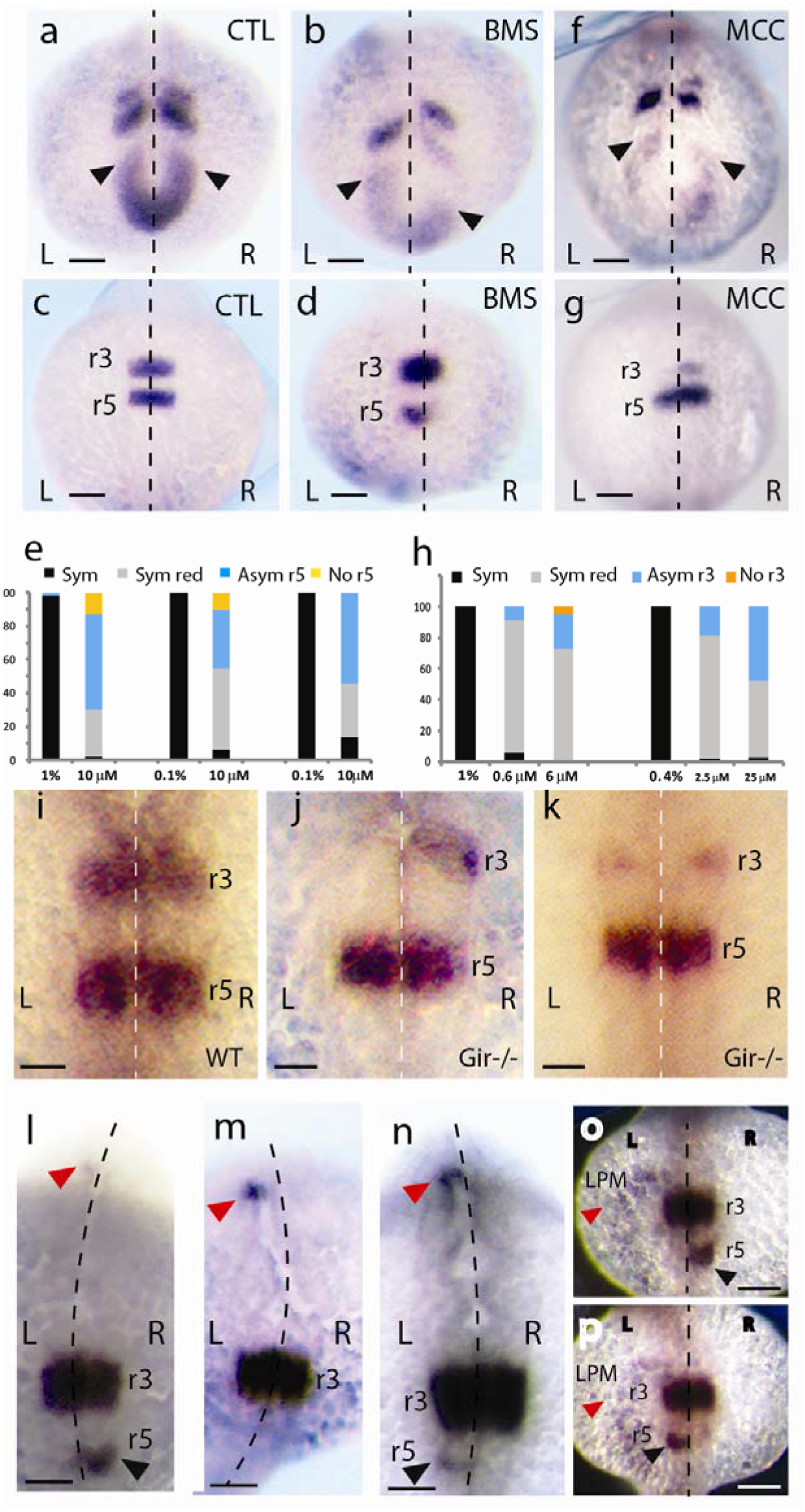
Inhibition and activation of RA signaling disrupts bilateral symmetry of the posterior hindbrain independently of Nodal signaling in zebrafish embryos. (a-d) zebrafish embryos control (a, c), or treated with BMS204493(BMS) (10 μM) (b, d) fixed at the 8-10 somite stage and hybridized with a *deltaC* (a-b) or a *krox20* probe (c-d). Arrowheads mark the tip of the traveling waves in the presomitic mesoderm. Scale bars: 100 μm (e) Percent of *krox20* bilateral symmetry phenotypes observed at 8 −10 somites following 10 μM BMS treatment at the 8-128 cell stage (right bars) compared to control treated with 1% or 0.1% DMSO (left black bars). Three different experiments are shown. Sym: symmetrical expression, Sym red: r5 expression reduced but symmetrical, Asymr5: any left-right asymmetric pattern of *krox20* in r5, No r5: r5 absent & r3-like stripe is enlarged due to anteriorization of the hindbrain (p = 0.0001 for for Sym. vs Asym) (f, g) Zebrafish embryos treated with MCC154 (MCC) (25 μM) fixed at the 8-10 somite stage and hybridized with a *deltaC* (f) or a *krox20* probe (g). Arrowheads mark the tip of the traveling waves in the presomitic mesoderm. Scale bars: 100 μm (h) Percent of *krox20* bilateral symmetry phenotypes observed at 8 −10 somites following treatment with various concentrations of MCC154 (MCC). Drug treatment (right bars) compared to control treated with 1% or 0.4% EtOH(left black bars). Two different experiments are shown: drug/vehicle addition at 50% epiboly (left side) or at the 128-cell stage (right). Sym: symmetrical expression, Sym red: r3 expression reduced but symmetrical, Asymr3: any left-right asymmetric pattern of *krox20* in r3, No r3: r3 absent (p < 0.004 for for Sym. vs Asym.) (i-k) *krox20* expression in 22-26 somites embryos from an incross of *giraffe^rw716^ cyp26a1* heterozygous mutants, (i) wild-type sib. (j) *gir-/-* homozygote with hemisegmental LR asymmetry of *krox 20* expression in r3. (k) *gir-/-* homozygote. Scale bars: μm (l-n) 24-25 somites zebrafish embryos treated with 10 μM BMS, hybridized with *krox20*and *lefty* probes. Red arrowhead shows *lefty1* expression in the diencephalon whereas black arrowhead in l and n shows asymmetric *krox20* expression in r5 (see Supplementary Table 3 for quantification). Scale bars: μm (o-p) 18-19 somites zebrafish embryos treated with 10 μM BMS,and hybridized with *krox20* and *southpaw* probes. Red arrowhead marks *southpaw* expression in the lateral plate mesoderm (LPM) whereas black arrowhead shows asymmetric *krox20* expression in r5. Scale bars: μm (a-g, i-p) Dorsal views. L: left, R: right. Anterior to the top.

Next, to probe the effect of RA excess, we analyzed *krox20* expression after inhibition of the Cyp26-class of cytochrome P450 enzymes, which are involved in the degradation of RA *in vivo*^21^. As observed with BMS, treatment with the Cyp26 inhibitor MCC154 led to desynchronized expression of *deltaC* between the left and right PSM (Fig. 1f). In the hindbrain, we also observed a striking phenotype where some MCC154-treated embryos exhibit LR asymmetric expression of *krox20*, which in extreme cases appears either on the left or the right side in r3 (Fig. 1g, Supplementary Table 2a). In contrast, the bilateral symmetry of the *krox20* r5 stripe was not significantly affected. In addition, there were classes of MCC154-treated embryos where *krox20* expression in r3 was reduced symmetrically or completely lost (data not shown and Supplementary Table 2a), in agreement with prior observations ^22,23^. These results were confirmed genetically by analyzing *krox20* expression in the *cyp26a1* mutant, *giraffe*^24^. 2% (5 of 318) of the embryos from *gir^rw716^* incrosses showed a LR asymmetric r3 phenotype for *krox20* and all of them genotyped as homozygous *gir* mutants (Fig. 1i-j, Supplementary table 2b). Another 2% showed phenotypes consistent with previous RA gain-of-function studies, namely a symmetric reduction of *krox20* in r3 ^22^ (Fig. 1k). All remaining embryos had symmetric *krox20* expression in r3 and r5 ^23^. The low penetrance of the asymmetric r3 phenotype in *cyp26a1* mutants suggests functional redundancy with *cyp26b1* and *cy26c1* and is consistent with the observation that the pan-Cyp26 inhibitor MCC154 generates asymmetric *krox20* r3 phenotypes at a much higher frequency (Fig. 1h, Supplementary Table 2a). Therefore, our data show that increasing or decreasing RA signaling disrupts both somite and hindbrain bilateral symmetry in zebrafish embryos.

Since in zebrafish, asymmetrical development of the brain-- leading to the LR specialization of the lateral habenula and parapineal gland--is controlled by the Nodal signaling pathway^25^, we next tested whether RA is involved in buffering asymmetric Nodal signaling. In zebrafish, the Nodal protein Southpaw is expressed in the left lateral plate and it acts in the left diencephalon via another Nodal protein, Cyclops to control LR asymmetry of the epithalamus ^25,26^. In BMS-treated embryos, there was no pronounced sidedness bias to the hindbrain *krox20* asymmetries [56% of cases with stronger disruption of *krox20* on the right side (176/ 316 embryos), 44% with stronger disruption on the left side (140/316 embryos)] (Supplementary Table 1 and data not shown). Expression of *lefty1*, which is induced by Cyclops in the left diencephalon, showed the same left expression bias in embryos treated with BMS and in control embryos (Fig. 1 l-n, Supplementary Table 3). Finally, we observed asymmetric r5 expression of *krox20* in the left or in the right side of BMS-treated embryos expressing *Southpaw* in the left lateral plate (Fig. 1o-p). Thus, the hindbrain asymmetries in RA-deficient embryos appear independent of the Nodal pathway.

The RA pathway involved in the control of somite symmetry requires the Rere chromatin-remodeling protein which together with Hdac1/2 and Wdr5, forms a coactivator complex called WHHERE ^4^. In zebrafish embryos homozygous mutant for the Rere homolog *rerea, babyface* (*bab^tb210^/ bab^tb210^*), normal somite symmetry was observed, possibly reflecting compensation by a maternal pool of *rerea* mRNA (Supplementary Table 3)^21^. Nevertheless, close to half of the homozygous mutant embryos (58 of 114 homozygotes) showed asymmetric pectoral fins at 5 days post-fertilization (dpf) (Fig.2a-b). Remarkably, a similar limb sidedness phenotype has been reported for several RA-pathway mouse mutants ^28,29^. In the hindbrain of *bab^tb210^/ bab^tb210^* embryos, we did not detect asymmetries of *krox20* expression (data not shown). We examined the distribution of reticulospinal neurons, which are hindbrain neurons relaying motor commands from the brain to the spinal cord and are arranged in a segmented, bilaterally symmetrical fashion ^30^. In *bab^tb210^/ bab^tb210^* embryos, all reticulospinal neurons were missing except for the Mauthner neuron in r4, which remained present and intact on both sides (Fig. 2c-d). However, in *bab^tb210^/+* embryos, we observed unilateral absence of one reticulospinal neuron in r6 in 43+/-1% of cases (n=37). The phenotype was observed either on the left or on the right side (Supplementary Table 4). The same phenotype was only observed in 15.5+/-1.5% of control wild type sibling (WT) embryos (n=13) (Fig.2c, e; Supplementary Table 4). Together, these data show that disruption of the RA coactivator Rerea function can result in LR asymmetries in a variety of embryonic tissues including limbs and hindbrain. However, Nodal-dependent epithalamic LR asymmetry, as evaluated with *lefty1*, appears normal in *rerea* mutants (Supplementary Table 3). This suggests that Rerea acts largely independently of Nodal to control bilateral symmetry of the developing zebrafish hindbrain.

**Figure 2:**
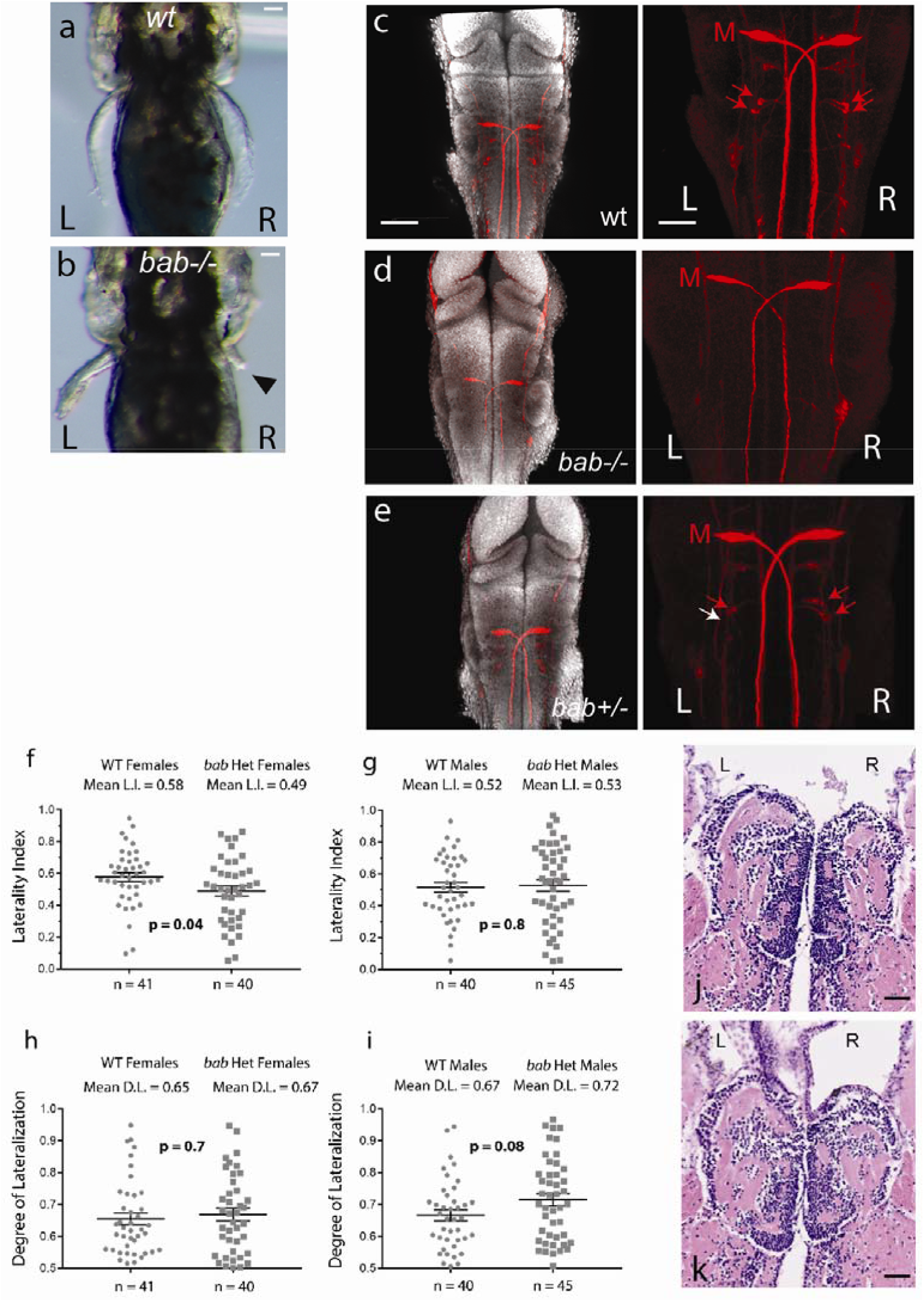
Laterality defects in the *rerea* RA co-activator zebrafish mutant *babyface* (*bab^tb210^*) (a, b) LR asymmetric fin (arrowhead) phenotype in *bab* homozygous mutant (b) compared to wild-type (WT) sib (a) in 8 dpf zebrafish larvae. Scale bar:50 μm (c-e) Dorsal confocal views of 48 hpf WT sib (+/+), *bab+/-* or *bab-/-* embryos showing rhodamine dextran-backfilled hindbrain reticulospinal neurons (red; left panels: merged view with DAPI counterstaining, right panels: red channel only). Note the asymmetry in the number of r6 reticulospinal neurons in *bab+/-* (red arrows point to reticulospinal neurons in r6, white arrow shows the position of missing neurons). M: Mauthner cell r4). Scale bars: left panels: 50 μm, right panels: 30 μm (f-g) Scatter Plots comparing population-level biases for clockwise swimming in WT adults vs *bab* heterozygotes, (f, g) Comparison of laterality indices (L.I.) between WT and *bab* heterozygotes females (f) and males (g). Numbers at top: group mean L.I. values. Horizontal black bars mark the mean L.I. and Standard Error of the Mean for the group of WTs or Heterozygotes. The clockwise swimming bias of females (f) is lost when one allele of *rerea* is disrupted (p = 0.04). For both WT and *bab* heterozygous males (g), no group-level directional swimming bias is evident. (h, i) Scatter Plots comparing the mean Degree of Lateralization (D.L.) between WT and *bab* heterozygotes. For both sexes, the p-value is > 0.05 for group level differences between WTs vs *bab* heterozygotes. Each gray square or circle represents the mean Laterality Index or Degree of Lateralization of one fish. (j, k) Swimming Laterality is independent of gross habenular Left-Right asymmetry. Coronal sections of 2 WT adult female brains. Although one fish (j) had a strong counter-clockwise swimming bias (90%) and the other (k) a strong clockwise bias (77%), the gross LR asymmetries of the dorsal habenular nuclei are the same (Left nucleus larger than Right). Scale bar: 50 μm (a-e) Dorsal views. L: left, R: right

As a functional output, we next tested whether *rerea* is required for normal, lateralized behavior in adult zebrafish. Since larvae homozygous for *rerea* mutations die around 8 days post-fertilization, we compared populations of WT adult fish with *bab^tb210^* heterozygotes (which are morphologically indistinguishable from wild type fish) for clockwise vs counter-clockwise swimming preference in circular swim tanks (Extended data Fig. 2 and Supplementary Movie 1). About forty *bab* heterozygotes and forty WT fish were assayed for each sex. Consistent with published reports ^31^, WT females showed a preference for clockwise swimming at the population level (mean Laterality Index, L.I. = 0.58; Fig. 2f). This population bias for clockwise swimming was lost in *bab* heterozygote females (mean L.I. = 0.49; p = 0.04; Fig. 2f). In contrast to published observations^31^, we saw no directional clockwise bias for WT and heterozygote males (Fig. 2g). We did not detect any change in the degree of lateralization (which measures the strength of the preference to swim in clockwise or counterclockwise direction) between *bab* heterozygote and WT females (Fig. 2h; p = 0.7) while a trend toward more lateralization was observed in male heterozygotes compared to WT (Fig. 2i; p= 0.08). Sex-specific differences in lateralized behaviors are common and may reflect interactions with sex-specific hormones and other factors ^32^. For both sexes, there was no statistically significant difference in swimming speed and swimming distance between *bab* heterozygotes and WT (Extended Data Fig. 3). Analysis of brain sections showed that gross LR asymmetry of the adult dorsal habenular nuclei was normal in a sampling of the *bab* heterozygote females that had been assayed for swimming laterality (9 of 9 fish, data not shown) and did not correlate with swimming direction. Moreover, strongly clockwise and strongly counter-clockwise WT fish exhibited the same habenular LR asymmetry (Fig. 2j-k). Collectively, these results argue that *rerea* modulates behavioral laterality through habenula- and Nodal-independent pathways. However, the limited understanding of zebrafish functional neuroanatomy precludes drawing any link between early hindbrain phenotypes and preferences in swimming orientation.

The data above prompted us to test whether the asymmetry-buffering role of RA discovered in the zebrafish brain extends to mammals. In the developing mouse cortex, a bilateral gradient of FGF signaling arising from the rostral patterning center (anterior neural ridge early and commissural plate later) plays a key role in specifying the different functional areas ^33^. High Fgf8 levels are required for motor cortex specification antero-medially whereas lower levels result in specification of sensory areas posterolaterally. This results in the establishment of a cortical map, which is strictly bilaterally symmetrical at this stage. Like in somites, this *Fgf8* gradient is also antagonized by RA, which acts to pattern the developing forebrain ^34^ in part through COUP-TFI (Nr2f1) which specifies the caudal part of the cortex including the sensory and visual areas ^35^. Consistently, an RARE-LacZ reporter^36^, which detects RA activity *in vivo*, is strongly expressed in the mouse developing forebrain and cortex (Fig. 3a) ^34,37^. When mouse null *Rere* mutants *openmind* (*Rere^om/om^*)^38^ were crossed to the *RARE-LacZ* reporter, the LacZ signal in the forebrain was strongly down-regulated ^7^. In addition, chromatin immuno-precipitation (ChIP) revealed that the Retinoic Acid Receptor alpha (Rara) and members of the WHHERE complex are bound to the RARE-LacZ promoter in the E13.5 mouse cortex (Fig. 3b-c). Finally, defects in forebrain and cortex development have been reported for RA-deficient mice and *Rere* mutants ^7,38–43^. Together, these observations suggest that Rere-dependent RA signaling plays a role in patterning the developing mouse cortex. Because *Rere^om/om^* mutants die at E10.5 before cortex formation ^38^, we analyzed cortical patterning in *Rere^+/om^* heterozygotes which are viable and fertile and which do not show any obvious behavioral difference with their WT littermates ^44^. Analysis of *Fgf8* expression using qPCR in dissected E10.5 brains showed an increase (^~^8%) in *Rere^+/om^* heterozygous mutants compared to WT sibling embryos, consistent with *Rere* antagonizing *Fgf8* expression as reported in homozygote mutants ^38^(Fig. 3d-f). At post-embryonic day 7 (P7), the motor cortex, whose boundaries are defined by the expression of *Lmo4*^45^, is expanded by ^~^10% in *Rere^+/om^* brains when compared to WT (Fig. 3g-i). A similar increase is observed in mutants deficient for RA signaling in the cortex^39^. In WT brains, the left and right motor cortices are of similar size, while in *Rere^+/om^* brains, the right motor cortex is ^~^5% larger than the left (Fig. 3g-h,j-l). We used magnetic resonance imaging (MRI) to identify cortical regions showing bilateral asymmetries of the gray matter in live WT and *Rere^+/om^* adult brains. The volume of gray matter in the motor cortex in *Rere^+/om^* was expanded asymmetrically on the right side compared to WT (Fig. 3m), consistent with the enlargement of the *Lmo4* motor domain observed in newborn (P7) *Rere^+/om^* brains. Together, these results demonstrate that *Rere*antagonizes*Fgf8* expression during cortical patterning. This also shows that *Rere* is involved in the control of motor cortex bilateral symmetry during development.

**Figure 3:**
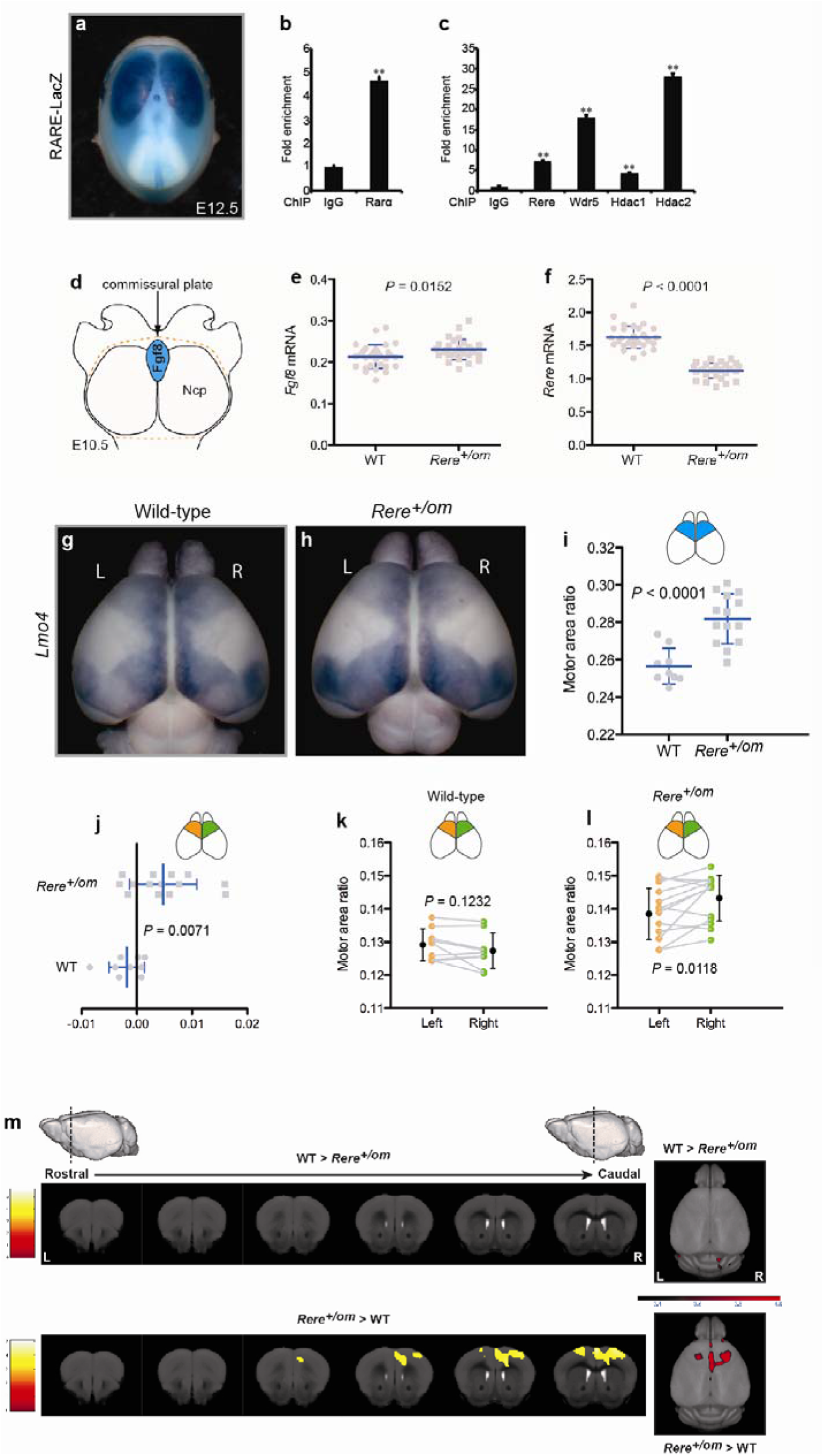
Retinoic acid signaling in the mouse cortex and asymmetric motor area expansion in newborn *Rere^+/om^* brains. (a) RARE-LacZ expression in the developing cortex of an E12.5 wild-type mouse embryo. Dorsal view, anterior to the top. (b, c) ChIP analysis of the *RARE* sequence on the RARE-LacZ reporter with specific antibodies against Rarα (b) or Rere, Wdr5, Hdac1 and Hdac2 (c) in E13.5 RARE-LacZ mouse cortices (n = 3). IgG: negative control. Data represent mean ± s.e.m. \*\**P* < 0.01. (d) Schematic dorsal view of dissected E10.5 forebrain showing the area of the neocortical primordia (Ncp) (orange dashed line, Anterior to the top) used for qPCR analysis of *Fgf8* (e) and *Rere* (f) expression from wild-type (n = 30) and *Rere^+/om^* (n = 29). Unpaired two-sample t-test, two-tailed, *Fgf8* mRNA: *P* = 0.0152 and *Rere* mRNA: *P* < 0.0001. (g, h) Dorsal views of wild-type (g) and *Rere^+/om^* (h) P7 brains hybridized *In situ* with a *Lmo4* probe. Left (L) and Right (R). Anterior to the top. (i) Motor area ratio (Area of rostral *Lmo4*-positive domain / Area of entire cortex) in wild-type (n = 9) and *Rere^+/om^* (n = 14) brains. Unpaired two-sample t-test, two-tailed, *P* < 0.0001. (j) Motor asymmetry index [(Right motor area – Left motor area) / Area of whole cortex)] in wild-type (n = 9) and *Rere^+/om^* (n = 14) brains. Unpaired two-sample t-test, two-tailed, *P* = 0.0071. (k-l) Left (Left motor area / Area of entire cortex) and Right (Right motor area / Area of entire cortex) motor area ratio comparison within the same brain in wild-type (n = 9) (k) and *Rere*^+/om^(n = 14) (l). Paired two-sample t-test, two-tailed, wild-type: *P* = 0.1232 and *Rere^+/om^*: *P* = 0.0118. In all graphs, data represent mean ± s.d. unless otherwise specified. (m) Magnetic resonance imaging from wild-type (n = 20) and *Rere^+/om^* (n = 18) adult brains. (Top) Coronal sections (left panels) and 3D brain projections (below background surface - max intensity) (right panel) representing regions larger in wild-type (WT >*Rere^+/om^*) (Bottom) coronal sections (left panels) and 3D brain projections (below background surface - max intensity) (right panels) representing regions larger in *Rere^+/om^* (*Rere^+/om^* > WT regions). The statistical parametric t-maps from Unpaired two-sample t-test are thresholded at *P* < 0.001 with an extent threshold of 500 voxels. The data are overlaid to an MRI template derived from Dartel normalization (using SPM8, Wellcome Trust Center for Neuroimaging, UK, http://www.fil.ion.ucl.ac.uk/spm) of all brain volumes (acquisition matrix size: 384×384, pixel size: 60×60 μm^2^; 72 slices, thickness: 0.22 mm). Yellow voxels indicate significant differences with the contralateral side. Increased volume of gray matter in the right hemisphere of *Rere^+/om^* mice by 4.6% compared to WT mice. The color bars represent t-scores. Left (L) and Right (R).

This prompted us to investigate whether *Rere* also controls the asymmetric localization of higher functions in the mammalian cortex. Handedness is a well-known lateralized behavior controlled by the motor cortex ^46^. Humans are the only species exhibiting close to 90% preferential bias toward usage of the right hand, a behavior controlled by the left hemisphere ^46^. In contrast, in WT mouse lines such as C57BL/6, around half of the individuals of a population show a consistent preference for the left paw (sinistral) while the other half exhibits a consistent preference for the right paw (dextral) when tested in specific behavioral assays ^47^. We analyzed forelimb usage in *Rere^+/om^* mice using an established assay of paw usage preference in adult mice (Mouse Reaching and Grasping - MoRaG, Supplementary movie 2)^48^. While ^~^35% (8 out of 24) of both WT females and males use their right paw to grab food pellets, in *Rere^+/om^* animals this proportion reached ^~^80% (19 out of 24) for both genders (Fig. 4a-d). No statistically significant difference in lateralization (defined as the consistency of paw usage within individuals) was observed between WT and *Rere^+/om^* (Fig. 4e-f). Overall, this analysis suggests that Rere controls brain functional asymmetry and hence the probability for an animal to use the right forelimb in a food reaching task at the population level.

**Figure 4:**
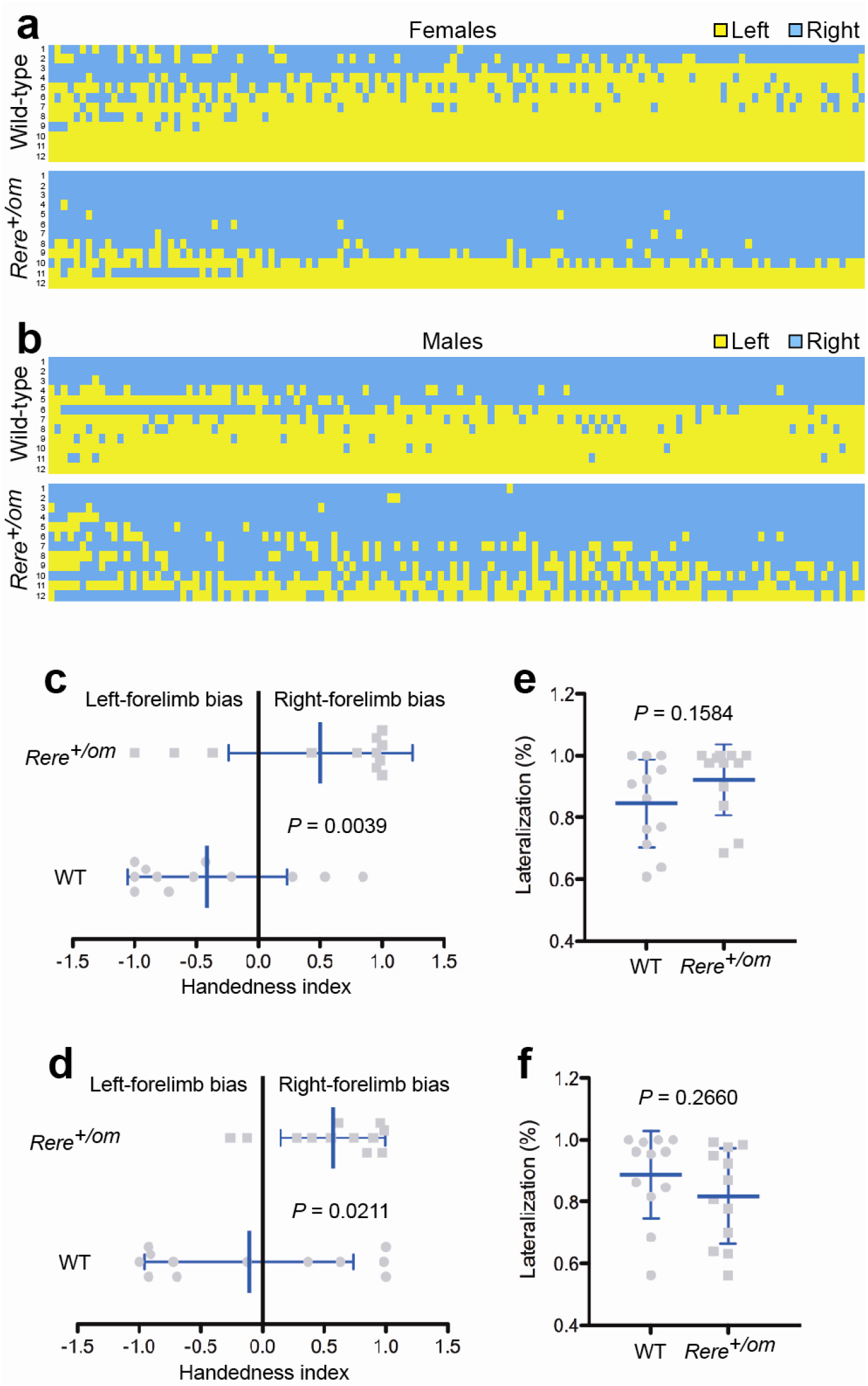
Rere controls asymmetric forelimb usage at the population level. (a-b) Heatmap detailing the mouse reaching and grasping (MORAG) behavior test in females (wild-type (WT): n = 12 and *Rere^+/om^*: n = 12) (a) and males (WT: n = 12 and *Rere^+/om^*: n = 12) (b). Columns correspond to 130 food pellets per animal trial. Yellow: Left-forelimb usage; Blue: Right-forelimb usage. (c-d) Handedness index [(Right-paw use - Left-paw use) / Total paw entries)] in WT and *Rere^+/om^* females (c) and in WT and *Rere^+/om^* males (d). Unpaired two-sample t-test, twotailed, females: *P* = 0.0039 and males: *P* = 0.0211. (e-f) Percentage of lateralization (highest value from either Right- or Left-forelimb use for each animal / Total paw entries) in wild-type and *Rere^+/om^* females (e) and in WT and *Rere^+/om^* males (f). Unpaired two-sample t-test, two-tailed, females: *P* = 0.1584 and males: *P* = 0.2660. In all graphs, data represent mean ± s.d.

This led us to explore the brain structure of adult *Rere^+/om^* animals. We used MRI to compare brain symmetry in a cohort of live WT and *Rere^+/om^* mice phenotyped for forelimb usage. Left and right comparison of the brains of sinistral and dextral WT mice by voxel-based morphometry (VBM) identified asymmetric differences in a region encompassing the right motor and somatosensory cortex (right sensorimotor cortex) only in animals using preferentially the right forelimb (Fig. 5a middle panels, c). This asymmetric sensorimotor region also included the cortex forelimb representation ^49^. No clear structural asymmetries were identified in the sensorimotor cortex of sinistral WT animals (Fig. 5a, top panels, c). A larger structurally asymmetrical region including the right sensorimotor cortex was identified in *Rere^+/om^* animals at the same position as the region identified in the brain of WT dextral animals (Fig. 5a, bottom panels, c).

**Figure 5.**
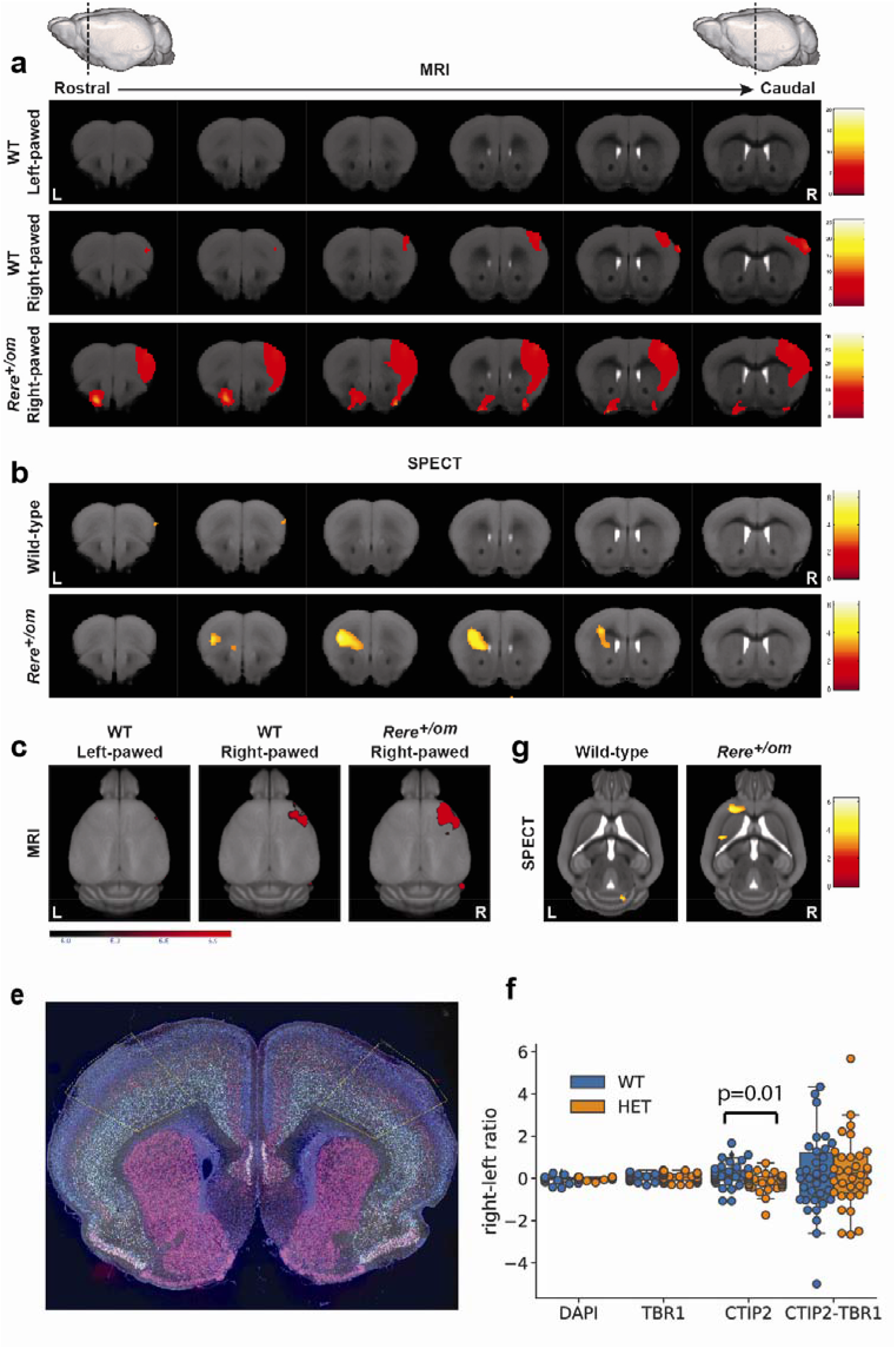
Asymmetric patterning of Corticospinal Motoneurons of the sensorimotor cortex in *Rere^+/om^* heterozygous mice. (a, c) Brain magnetic resonance imaging from wild-type (WT) sinistral (n = 12), WT dextral (n = 8) and *Rere^+/om^* dextral (n = 18) animals. Coronal sections (a) and 3D brain projections (below background surface - max intensity) (c) representing structural Left-Right asymmetry differences. Increased gray matter in the right hemisphere of dextral WT and dextral *Rere^+/om^* mice by 6% and 7.5% respectively compared to the left hemisphere. The statistical parametric t-maps from uncorrected Paired two-sample *t*-test are thresholded at *P* < 0.00001 with an extent threshold of 500 voxels and overlaid to the MRI template. (b, g) Single photon emission computed tomography (SPECT) in wild-type (n = 9) and *Rere^+/om^* (n = 9) animals. Coronal (b) and transverse (g) sections representing Left-Right asymmetry differences in regional cerebral blood flow. The statistical parametric t-maps from One-sample t-test for Left-Right asymmetry are thresholded at *P* < 0.01 and overlaid to the MRI template. The color bars represent t-scores. Left (L) and Right (R). (e) Coronal section of the brain of a P7 *Rere^+/om^* mouse at the level of the sensorimotor cortex labeled with antibodies against Ctip2 (purple) and Tbr1 (green) which respectively label the layer V and VI neurons. Orange lines delimits the area where cell counts were performed. (f) Comparison of the left-right ratio in the whole cell population and in Ctip2, Tbr1 and double positive populations of neurons counted in the orange boxes shown in (e) between WT and *Rere^+/om^* mice (HET). The only difference detected is for the Ctip2 positive cells (p=0.01, Mann-Whitney test) which include the corticospinal motoneurons of Layer V.

The apparent increase in the right motor cortex volume in *Rere*^+/om^and dextral animals is unexpected as the right paw-preference is controlled by the left hemisphere. In cortices of mice mutant for *COUP-TFI*, the motor cortex expands into the sensory area resulting in improper patterning of the cortico-spinal motor neurons (CSMN) in the ectopically motorized area ^50^. This leads to an impairment of fine motor skills in mutant mice. The slight caudal expansion of the right motor cortex in *Rere^+/om^* mice could similarly lead to inappropriate specification of a subset of CSMN in the left forelimb projection area thus resulting in decreased dexterity associated to the left paw. To investigate CSMN specification in *Rere^+/om^* mutant brains, we analyzed the expression of Ctip2 (which strongly labels the CSMN of layer V) and Tbrl (which is expressed by layer VI neurons), in sections of the motor cortex region showing asymmetries in mouse P7 brains. We found that compared to WT, *Rere^+/om^* mutants show similar total numbers of neurons, of Tbr1 neurons, and of Ctip2-Tbr1 double-positive neurons on the left and right side, but a slight yet significant decrease in Ctip2-positive neurons in the right side (Extended data Figure 4). Thus, our data show subtle abnormal LR patterning of the CSMN neurons in the right side of *Rere^+/om^* mutant brains which could account for the right dominance of the heterozygous mutant population.

We next performed brain perfusion Single Photon Emission Computed Tomography (SPECT) to analyze regional cerebral blood flow as an overall indicator of neuronal activity. In *Rere^+/om^* but not in WT mice, we observed asymmetric tracer uptake in a deep region including part of the motor and somatosensory cortex in the left hemisphere (Fig. 5b,g). Interestingly, the tracer uptake signal was approximately at the same antero-posterior level of the sensorimotor cortex (albeit on the contralateral side) as the structural asymmetry detected by MRI (Fig. 5a,c). This data suggests increased brain activity in the left hemisphere of *Rere^+/om^* brains, consistent with their preference in using the right forelimb.

Taken together, our work shows that the level of Rere-dependent RA signaling controls the degree of brain asymmetry at the morphological and functional levels in zebrafish and mouse embryos. Our data suggest that this pathway controls the LR symmetry of the FGF signaling gradients involved in brain patterning. In zebrafish, FGF signaling acts as a left determinant^10^ which also controls aspects of epithalamus asymmetry ^51,52^. In humans, SETDB2 (which antagonizes FGF signaling to control visceral asymmetry in fish) has been implicated in the control of handedness^9,53–55^. Our studies show that altering RA levels can generate LR asymmetries in the zebrafish hindbrain independently of Nodal. In mammals, evidence for implication of Nodal signaling in brain asymmetry is very limited ^53^. The frequency of left-handed individuals is not increased in patients with *situs inversus totalis* arguing against a role for this pathway in the control of handedness ^56^. Together, these data suggest that the Rere-dependent RA pathway controls brain asymmetry independently of Nodal.

Our study demonstrates that reduction of the dosage of the RA-coactivator Rere can dramatically shift handedness to the right at the population level in mouse. While handedness has long been known to exhibit a genetic component, its transmission follows a complex non-Mendelian or polygenic mechanism which is not understood ^57^. A classical model of genetic inheritance of handedness called the right shift theory proposed that handedness is defined stochastically, resulting in the roughly equivalent distribution of dextral and sinistral individuals as observed in most animal species ^58^. In humans however, a right shift gene would skew the distribution to the right, leading to the strong right bias observed. Our data show that *Rere* behaves as such a right-shift gene. Genetically, RA levels can be modulated in many different ways, which could account for a multifactorial genetic control of handedness. The ability of RA levels to affect bilateral symmetry of the hindbrain region observed in zebrafish might also be relevant to handedness as recent studies argue that hindbrain and spinal cord asymmetries may be the earliest steps in the mechanism establishing human handedness ^59,60^. *RERE* has been identified in GWAS studies as a susceptibility locus for schizophrenia ^61^. This neurological disease is associated with brain symmetry defects and increased numbers of left-handed and mixed-handedness individuals suggesting that RERE and the RA signaling pathway could be involved in their etiology ^62^. Thus, our study provides a basis to understand the lateralization of behaviors such as handedness in the mammalian cortex. This mechanism might also apply to the asymmetric distribution of higher cognitive functions such as speech in the human brain.

## AUTHOR CONTRIBUTIONS

### Authors’ contributions

M.R. and G.C.V.-N respectively designed, performed and analyzed the zebrafish and mouse experiments with O.P. M.R. performed the zebrafish embryo treatments and mutant production and analysis with help from J.V. M.S.G. and C.S. prepared the CYP26 inhibitor MCC154. M.L., L. B-C and M. R. performed the behavioral tests. D.C., S.B. and N.D. performed the analysis of reticulospinal neurons. J.- L.P. and G.C.V.-N. performed the qPCR and qChIP experiments. G.C.V.-N. did the mouse embryo and brain analysis. A.M. performed the neuron quantifications. F.R. and H.M. performed the MoRaG test. A.P. and S.L. designed, performed and analyzed the MRI study. P.L. and D.B. performed the SPECT experiments. V.N. performed the SPECT image analysis. G.C.V.-N. designed and analyzed the MoRaG, MRI and SPECT experiments and results. M. R., G.C.V.-N. and O.P. wrote the manuscript and OP supervised the project.

## Acknowledgements

We thank members of the Pourquié laboratory and C. Tabin, F. Guillemot, D. Henrique and A. Chedotal for critical reading and comments on the manuscript. We are grateful to the IGBMC zebrafish core facilities and to Dr. S. Vincent for assistance with some of the statistical analysis and to Virgile Bekaert et Ali Ouadi for help with the SPECT experiments. We thank N. Plaster, T. Schilling and the Zebrafish National BioResource Project, Japan, for providing fish lines and A. Peterson, and J.

Rossant for providing mouse lines. Research in the Pourquie lab was supported by an advanced grant of the European Research Council to OP. We thank R. Valabrègue from CENIR for his help in preprocessing the MRI data. The research leading to the MRI results has received funding from the programs “Investissements d’avenir” ANR-10-IAIHU-06 and “Infrastructure d’Avenir en Biologie Santé” ANR-11-INBS-0006. Work in the Bally-Cuif lab was supported by the European Research Council (AdG322936) and Labex Revive. Mohamed Sayef was supported by a grant from Cancer Research UK (Grant Ref. C7735/A9612)

## METHODS

### Zebrafish Lines

The *giraffe^rw716^* line ^24^ was obtained from the Zebrafish National BioResource Project, Japan. The *bab^tb210^* allele^27^ was obtained courtesy of Nikki Plaster and Tom Schilling. Zebrafish embryos and larvae were staged in cell numbers, somite numbers and in hours post-fertilization (hpf) or days post-fertilization (dpf).

### Drug Treatments

BMS-204493 (BMS), a pan-RAR antagonist, was custom-synthesized and dissolved in DMSO to a stock concentration of 10mM. Unless otherwise noted, experiments were done using 1-10 μM BMS, a dose range that specifically inhibits RA signaling in zebrafish embryos^15^. The pan-Cyp26 inhibitor MCC154 (3-imidazol-1-yl-2-methyl-3-[4-(naphthalen-2-ylamino)-phenyl]-propionic acid methyl ester) was synthesized as described ^63^ and resuspended in ethanol (6mM stock). All drug treatments used vehicle alone (DMSO or EtOH) as the negative control. BMS was added at the 8-128 cell stage (unless otherwise noted) in fish system water and embryos were incubated in BMS at 28.5°C until the time of analysis. For a few experiments, development was slowed down by incubation at 23.5C°C. MCC154 was added at either 3 hpf or 6 hpf, at doses from 0.25 to 64 μM. To inhibit pigment formation, PTU (1-phenyl 2-thiourea) was added at 10hpf (bud stage) to a final concentration of 0.003%. Fish used in the swimming behavior assay were never exposed to PTU during their lifetimes.

### Zebrafish *in situ* hybridization, histology and reticulospinal neurons retrograde labeling

Whole mount *in situs* with digoxygenin-labelled RNA probes and development with BMPurple (Roche) were performed as described previously ^64^. *krox 20* probe, *pBluescript KS+Krox 20* (Sal I digest; T7 Pol); southpaw probe, *pGEMT-Southpaw1.4* (Spe I; T7 Pol); *delta C* probe, *pL3-Delta C* (Xba I; T7 Pol); *unxc4.1* probe (EcoRV; SP6 Pol); *lefty1* probe; *pAD-Gal4Lefty1* (Mlu I; T7 Pol); *hoxb1a* probe, *pBluescript KS+hoxb1a* (Sal I; T7 Pol) (clone ibd3532).

Adult brains sections were prepared as follows: adult zebrafish were euthanized; the heads were cut off with razor blades and fixed in 4% PFA for 24-48 hours. Brains were dissected and stored in 1XPBS + azide until use. Brains were dehydrated and embedded in paraffin, sectioned at 7 μM and stained with hematoxylin-eosin which gave better contrast of the habenulae than the standard Nissl staining.

<u>Retrograde labeling</u>: reticulospinal neurons were backfilled by applying a 5% solution of 10,000 MW Tetramethylrhodamine Dextran (Thermo Fischer) at the level of a spinal cord transection posterior to the hindbrain in 48hpf live embryos. The dye was allowed to diffuse for 6 hours, after which time the embryos were fixed, incubated in DAPI and processed for whole-mount confocal scanning.

### Genotyping

The following combinations of genomic PCR primers and restriction digests were used for genotyping zebrafish adults or embryos: *bab^tb210^* allele (mutation causes the loss of an RsaI site within the PCR amplified region)^27^: REREAExtFwd: 5’-GTATATGTAGTTCTTGATGTCAGTTGTTATGGG-3’ REREAExtRev: 5’-GTGATTCCGTACCGAAGTTAAAGTTTTGTGC-3’*gir^rw716^* allele (mutation creates an XbaI site within the PCR amplified region): cyp26AlGFl: 5’-CAGGGTTTGAGGGCACGCAATTT-3’ cyp26AlGRl: 5’-G CTGCTTCTTTCATCGCCTAAGC-3’ After *in situ* hybridization and color development of embryos, genomic DNA was extracted from the embryos using the following protocol (method from M. Halpern lab, personal communication): 1X Extraction Buffer: 1.5 mM MgCl_2_,10 mM Tris HCI pH 8.3, 50 mM KCl, 0.3% Tween 20, 0.3% NP-40. Single embryos were rinsed with 1X PBS. 50 μl of extraction buffer was added to each embryo, heated to 98°C to 100°C for 10 minutes and then incubated for 2.5 hours at 55°C with Proteinase K added to a final concentration of 2 mg/ml. After heating at 98°C for 10 minutes, the extracts were centrifuged briefly, and the supernatants were transferred to new tubes and stored at - 20°C until use. PCRs were carried out in 50 μl reactions using 1X Promega GoTaq Green Flexi Buffer, 1 ng/nl of each primer, 2 mM MgCl_2_, 200 μM dNTPs, 5 U Taq and 2 to 10 μl of extract. Genomic PCR reaction conditions were 92°C, 2 min followed by 44 cycles of 95°C, 60s, 65°C, 30s, 73°C, 60s then 4°C hold.

### Laterality Behavioral Assay

Assays were done at the CNRS AMATRACE Behavior Core Facility in the Zebrafish Neurogenetics Lab at Gif-sur-Yvette (France). Zebrafish embryos from outcrosses of several *bab^tb210^* heterozygotes (in the AB background) to AB fish were raised in 40-liter tanks under defined day-night conditions and sorted at approximately 9 months of age into wild-type (+/+) and *bab* heterozygote fish. Because we planned to use the adults in swimming assays, we decided not to genotype the fish by PCR on DNA from fin clips. This was to avoid any confounding variables due to variable fin regeneration and trauma. Consequently, the genotype of each fish was verified by two ID crosses, scoring for the presence of at least two out of three visible phenotypes in the progeny (reduction of pectoral fins, outgrowth of the Retinal Pigmented Epithelium (RPE) and jaw defects), which would indicate the presence of homozygous *bab^tb210^* mutant embryos and of heterozygous parents. Diseased and non-thriving fish were discarded. The sex of each fish used in the laterality assay was determined by visual criteria and in ambiguous cases, by retrospective dissection of fish. “Donut”-shaped swimming chambers were used for assays of rotational swimming behavior and were made by immersing a smaller plastic bucket (15 cm diameter) into the center of a larger circular plastic tank that was infrared-transparent [KIS Ecobowl, 36 cm (bottom diameter) X 38 cm (top diameter) X 18 cm (height), catalog #008712, polyethylene]. This gave a “donut”-shaped swimming chamber with an opaque white inner wall and a semi-translucent blue outer wall and floor. Both inner and outer walls were partially reflective. Tanks were filled to a depth of 7.5 cm prior to each assay. Each individual adult was transferred by dip net into a swimming chamber. Videos of fish in one to five swimming chambers were then recorded simultaneously using Viewpoint Life Sciences Infrared Cameras and Platforms with Zebralab tracking software, and the number of counter-clockwise (CCW) and clockwise (CW) rotations for each fish was counted with Viewpoint’s automated rotation counting software. Program parameters for the rotation counting software were set to: minimum diameter 6.0 cm, return angle 40.0 degrees, scale calibrated to the water surface level. In any cases where infrared tracking was imprecise, the videos were played back, and the number of clockwise and counterclockwise rotations was scored visually. To avoid introducing a bias, if the number of visually scored replacement videos was different between the WT and heterozygote samples, we replaced additional rotation scores counted by the automated software with visually determined rotation scores, so that the total number of visually scored assays was the same for both the WT and the heterozygous sets. For visual scoring, complete rotations were defined as either continuous 320 - 360 degrees swims or circular swims that were briefly interrupted by a pause in swimming or by a brief reversal of swimming direction. The accuracy of the automated rotation counting software was cross-checked by comparing the scores obtained by automated and visual counts of rotations for twenty individual assays (25 minutes recording periods). Each fish was recorded for twenty-five minutes and the assay was repeated on 3 to 4 successive days. Fish often showed reproducible directional preferences from one day to the next (Extended Data File Fig. 2c and Supplementary Tables 5, 6). For each fish, this resulted in 3 to 4 daily laterality indices, which then were averaged to give a mean laterality index (L.I.) for each fish, which were in turn averaged to get a mean L.I. for the WT or heterozygote group. The laterality index, L.I., is defined as the ratio of the number of complete CW rotations/swims divided by the sum of *ON + CON* rotations. L.I. < 0.5 indicates a preference for swimming in the counter-clockwise direction. Laterality indices were measured for about 40 wild-type females and 40 *bab^tb210^* heterozygous females and for a similar number of WT and heterozygous males. Mean laterality indices and rotation scores for all tested fish are listed in Supplementary Tables 5 and 6.

We also calculated the Degree of Lateralization, D.L., which is independent of direction. Thus, a fish which completes 90 clockwise rotations and 10 counter-clockwise rotations during the assay period has the same D.L. index (0.9) as a fish that completes 90 counter-clockwise rotations and 10 clockwise. Mean D.L. indices for the populations were obtained by calculating a mean D.L. index for each fish and then obtaining the population mean D.L from those (Supplementary Table 7 and 8).

Locomotion parameters during the rotational assays (Extended data figure 3) were extracted using the Zebralab tracking software. We plotted the speed and distance travelled for all fish tested in the rotational assays except for a few of the males *bab* Hets due to some minor tracking failure.

#### Statistical Analysis

For each population of fish (wild-types or *bab* heterozygotes of the same sex), the distribution of mean laterality indices followed a normal distribution, as verified using the Shapiro Wilk and D’Agostino-Pearson omnibus tests. Since Fisher tests indicated the variances were similar (homoscedastic), an unpaired, two-tailed t-test was used to compare the mean laterality indices of the two populations (WT vs heterozygotes of the same sex) and calculate p values. For other parameters (D.L., swimming speed and swimming distance), group means were compared, and p-values determined using the Mann-Whitney test.

### Mice breeding and generation of mutant embryos

The generation of the *Rere^+/om^* mouse line was described previously^38^ and the line was maintained on a C57BL/6 genetic background. To study the status of retinoic (RA) signaling, the *RARE-LacZ* reporter mice,^36^ in which LacZ expression is driven by the Retinoic Acid Responsive Element (RARE) of the retinoic acid receptor beta gene (*Rarβ*), were crossed to WT C57BL/6 and the resulting embryos were analyzed for *6-*galactosidase activity using X-gal (5-bromo-4-chloro-3-indolyl-β-D-galactopyranoside). Embryos were stages in embryonic days (E).

### Mouse embryo and brain whole-mount *in situ* hybridization

*In situ* hybridizations were performed as described ^65^. The probes used are described in the literature: *Uncx4.1*^66^, *Lfng*^67^ and *Hes7*^68^. To generate the *Lmo4* probe, the full-length cDNA encoding mouse *Lmo4* was cloned into pCMV-Tag2.

### Real-Time PCR Assays using NIH3T3 cells and dissected neocortical primordium/commissural plate from mouse embryo brain

Total RNA was isolated from NIH3T3 cells using the FastLane Cell One-Step Buffer Set (Qiagen). The mouse embryo brains at E10.5 were dissected in cold PBS. Only the neocortical primordium and the commissural plate were dissected and stored in RNA Later (Qiagen) at −20°C until RNA extraction. The total RNA was purified using Qiazol and the RNeasy Mini Kit (Qiagen). All the qPCR experiments were performed using Quantifast SYBR Green RT-PCR Kit (Qiagen) in a LightCycler 480 II System (Roche). The following primers from Qiagen (Qiagen Quantitect Primer Assay) were used:

| Gene | Catalogue number |
| --- | --- |
| *Rere* | QT01075774 |
| *Rplp0* | QT00249375 |

For *Fgf8* expression analysis from dissected mouse brains the following primer pair was used:

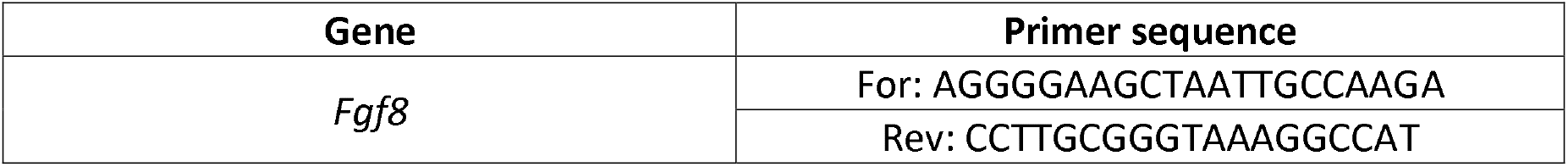

### Chromatin Immunoprecipitation (ChIP) Assay

For ChIP assay with mouse embryonic tissue or brain cortices, E8.75-9.0 embryos or E12.5-13.5 brains were harvested from transgenic *RARE-LacZ* mice and 20 μg of chromatin were used per immunoprecipitation. ChIP was performed as described ^7^. Antibodies used for ChIP are as follows:

| Antibody | Catalogue number (Company) |
| --- | --- |
| Rere H-113 | sc-98415 (Santa Cruz) |
| Wdr5 | 07-706 (Millipore) |
| Hdac1 C-19 | sc-6298 (Santa Cruz) |
| Hdac2 H-54 | sc-7899 (Santa Cruz) |
| Rarα C-20 | sc-551 (Santa Cruz) |

qPCR was performed with LightCycler 480 SYBR Green I Master mix (Roche) in a LightCycler 480 II System (Roche) using the following primers:

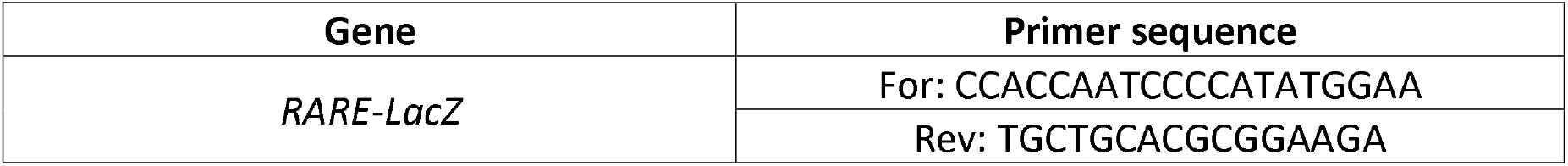

### Mouse Reaching and Grasping (MoRaG) behavior test

The MoRaG test consists of an assessment of reaching and grasping motor behavior in mice. These specific fine-tuned motor abilities are altered in neurological diseases such as Parkinson’s disease. In this test, mice submitted to food restriction are forced to use a single paw instead of their mouth in order to get a small food pellet. Mice are tested in the Collin’s apparatus ^69^, which consists of a Plexiglas chamber with dimensions of ^~^10 cm high, by 6 cm deep and by 6 cm wide. On the outside of the front wall of each cubicle, a Plexiglas feeding platform, accessible only through a ^~^10 mm diameter opening, is attached ^~^5 cm from the floor. A total of 24 wild-type (12 females and 12 males) and 24 *Rere^+/om^* (12 females and 12 males) were used for the MoRaG test to identify the preferred forelimb used to retrieve the food pellet. Mice are food-deprived overnight in order to increase their motivation. On the day of the test, the mouse is placed in the testing box and allowed to acclimate for 5 minutes before the task begins. A small food pellet is placed on the feeding platform. It can be accessed through the opening but the size of the opening is not large enough to allow the mouse to collect it by using its mouth. The MoRaG performance scale is designed to record the upper-forelimb movement for reaching (the forepaw proceeds in the horizontal plane out of the trunk to approach an object at a distance) and grasping (the object is grasped). The experimental procedure comprises 3 testing sessions. On the first session it was done for 30 trials and on the next 2 sessions 50 trials each were done for a total of 130 trials, and accuracy (percentage of success) of reaching and grasping was recorded. The motor lateralization was also evaluated to analyse the paw of preference to reach the food pellet. The behavioral pipeline was performed in agreement with the EC directive 2010/63/UE86/609/CEE and was approved by the local animal care, use and ethics committee of the IGBMC (Com’Eth) under accreditation number (2012-139).

### Magnetic Resonance Imaging (MRI)

#### 1. MRI data acquisition

A total of 22 wild-type and 22 *Rere^+/om^* mice were imaged after performing the MoRaG test. All images were acquired using a Bruker Biospec 117/16 USR MRI system (BGA-9S gradients, 750 mT/m, AVIII) running Paravision 5.1. A 72 mm resonator was used for signal emission and a planar surface coil for mouse heads was used for signal reception (Bruker Biospin, Ettlingen, Germany). Structural T2-weighted images were acquired for all mice. A two-dimensional turbo-RARE (Rapid Acquisition with Relaxation Enhancement) sequence was used with TR = 6500 ms; TE= 40 ms; matrix = 384 x 384; field-of-view (FOV) = 23 x 23 mm^2^ (60 x 60 μm^2^ in-plane resolution); 72 slices; slice thickness = 0.22 mm; number of averages = 4; scan time = 2h 8min. The animals were anesthetized using Isoflurane at 1.5-2% mixed in oxygen:air (1:5). They were placed in a mouse cradle and their heads were held fixed with ear bars and a tooth bar. A water-circulating system integrated to the cradle ensured that the animals were kept at physiological temperature. Their respiration was monitored via a pressure pad placed under their abdomen. Physiological parameters were monitored throughout the MRI sessions.

#### 2. MRI data analysis

##### 2.1. Voxel-based morphometry and preprocessing

We first multiplied the voxel size by a factor of 17 in order to have brain dimensions similar to human brains to match the default mode parameter of Statistical Parametric Mapping software (SPM8, Wellcome Trust Centre for Neuroimaging, UK, http://www.fil.ion.ucl.ac.uk/spm). Image segmentation and normalization was performed in two steps using first the template given by SPMMouse and then a symmetrical template. First, SPM8 was used to segment into gray matter (GM) and white matter (WM) probability maps and to normalize the structural images with the segment function and the template given by SPMMouse ^70^. Dartel normalization was performed using the GM and WM probability maps. Dartel normalization was performed on all structural and swapped images (to invert left and right sides), which were then averaged. This provided a symmetrical template. Second, we performed again the segmentation and Dartel normalization on native images using the symmetrical template. The resulting normalized modulated GM images were then smoothed with an 8 mm kernel before performing voxel-based morphometry (VBM) analysis.

##### 2.2. Statistical analysis

The left and right hemispheres were considered for comparisons. Statistical analyses were conducted to assess within-group differences in GM between the left and the right hemispheres by comparing native and swapped images. Statistical analyses were also conducted to compare wild type and *Rere^+/om^* volumes for between-group comparisons. Both Paired two-sample t-test for within-group statistical analyses and Unpaired two-sample t-test for between-group statistical analyses were run in SPM8. Clusters were considered significant at P < 0.05 with small volume correction (SVC, sphere radius = 3 mm, i.e. 3 voxels radius) with an uncorrected height threshold of *P* < 0.00001 for within-group comparisons and *P* < 0.001 for between-group comparisons, and an extent threshold of 500 voxels.

##### 2.3. 3D brain projections

For 3D brain projections (below background surface - max intensity) to compare wild type versus *Rere^+/om^* and Left versus Right asymmetric differences, the software used for volume rendering was from MRIcron image viewer (Neuroimaging Informatics Tools and Resources Clearinghouse, NITRC, http://www.nitrc.org/projects/mricron, 2013 release).

### Single photon emission computed tomography/Computed tomography (SPECT/CT)

#### 1. SPECT/CT data acquisition

A total of 9 wild-type and 9 *Rere^+/om^* mice were imaged to detect regional cerebral blood flow. In order to recognize the different organs, prior to SPECT analysis, a CT image was recorded on the AMISSA platform, a homemade multimodality imaging system for small animals combining X-ray, SPECT and PET devices. The μCT delivered a 3D reconstructed volume of the animal in real time ^71,72^. Targeting of 99mTc-HMPAO was monitored via the μSPECT imaging technique. Briefly Technetium-99m, as [^99m^TcO_4_] Na in physiological solution, was obtained from a ^99^Mo/^99m^Tc generator Elumatic^®^ III (CisBio/IBA Molecular, France). The [^99m^Tc]-HMPAO was prepared using the commercial kit Cerestab (GE Healthcare, Vélizy, France) following provider’s recommendations for preparation and quality control. Mice were anesthetized by intraperitoneal injection of 10 μl.g ^-1^ of a solution made with ketamine hydrochloride 10% (Imalgene, Centravet, Velaine en Haye, France) and Xylazine 5% (Rompun, Centravet, Velaine en Haye, France) and placed in prone position in the animal holder after injection in the tail vein. μSPECT imaging was performed 4 minutes post-intravenous injection of 30 MBq (in mean) of 99mTc-HMPAO injected in the tail vein. The μSPECT system consists of a four heads detection gamma camera. Each head comprises five-separated detection modules arranged along a circle of 58 mm with the pinhole as the center. A detection module consists of a YAP:Ce matrix of 8 x 8 scintillating crystals 2.3 x 2.3 x 28 mm^3^ each coupled to a multi-anode photomultiplier 8 x 8 (Hamamatsu H 8804). The distance from the pinhole to the axis of rotation is 28 mm and the distance between the pinhole and the crystal is 58 mm, which results in a magnification factor of 2.07. Images were reconstructed using the OSEM (Ordered Subset Expectation Maximisation) iterative algorithm adapted for pinhole imaging. Images were viewed and quantified using the Anatomist freeware (http://brainvisa.info/index_f.html).

#### 2. SPECT image analysis

SPECT image analysis is conducted according to the four following steps: (i) brain extraction, (ii) asymmetry map calculation, (iii) registration in a common space and (iv) voxel-based statistical analysis. The first three steps are performed using medipy software (https://piiv.u-strasbg.fr/traitement-images/medipy/). Voxel-based statistical analysis is conducted using SPM (http://www.fil.ion.ucl.ac.uk/spm/).

##### 2.1. Brain extraction

The first step consists in extracting the brain from the whole-body SPECT acquisition. To this end, we use the anatomical information carried out by the CT image, which is in the same physical space as the SPECT image. The skull is segmented from the CT image by using an intensity thresholding technique (Otsu’s method) and by affinely registering a probabilistic prior to remove bones that do not belong to the skull. Then, a mask of the brain is built from the skull using morphological operators dedicated to cavity filling. This brain mask is then applied on the SPECT image to extract the brain perfusion map.

##### 2.2. Asymmetry map calculation

The inter-hemispheric symmetry plane is then estimated from each brain perfusion map by minimizing the squared intensity differences between the original and the swapped images. Then, an asymmetry map is computed as the difference between the original and swapped images divided by the sum of the original and swapped images. By this way, the values of the asymmetry map range from −1 to 1 and are made independent from the perfusion amplitude.

##### 2.3. Registration in a common space

To conduct voxelwise analysis of these asymmetry maps across a population, they should be registered in a common space. This is done by affinely registering each SPECT image on a study specific template, and by applying the estimated transformation to the corresponding asymmetry map. The study specific template is built from both wild-type and *Rere^+/om^* images. This is done by affinely registering all SPECT images on an arbitrary chosen image. Then, an average image is computed from all registered images. Then, all images are registered again on the average image and a new average image is built. This is done until the process converges to obtain the final template.

##### 2.4. Voxel-based statistical analysis

All registered asymmetry maps are smoothed using a Gaussian kernel (FWHM: 1 mm) and a One-sample *t*-test is conducted voxelwise to find out the areas where the brain perfusion is asymmetric within each group. Statistical maps are thresholded at *P* < 0.01 (uncorrected P-value).

### Histology and quantification of neuronal populations in mouse brain sections

P7 mouse brains were perfused with 4% paraformaldehyde, dehydrated in ethanol and embedded in paraffin prior to sectioning. 7 μm were obtained and subsequently processed for immunohistochemistry with anti-Tbr1 and anti-CTIP2 antibodies. The asymmetric region of the sensorimotor cortex of dextral animals identified by MRI was manually drawn on one hemisphere using the image analysis software Fiji^73^. The symmetric region was then drawn on the other hemisphere and both regions were cropped and analyzed independently. Nuclei were automatically segmented on each channel using the image analysis software CellProfiler^74^. In short, the intensity of each channel was normalized and the segmentation threshold to distinguish positive and negative nuclei was manually set for each channel but kept constant for every brain slice. The number of positive nuclei per channel was counted for each hemisphere. The number of single and double-positive nuclei (CTIP2: Red+, TBR1: Green÷) and the total number of cells labeled with Hoechst was measured. The normalized excess between the two hemispheres ((right-left)/left) was then calculated for all the brain slices. Similarly, the area difference between the two sides was calculated using the same index. The significance of the difference between the wild type and *Rere^om^* heterozygotes excesses was assessed using the Mann-Whitney test.

**Extended data Figure 1:**
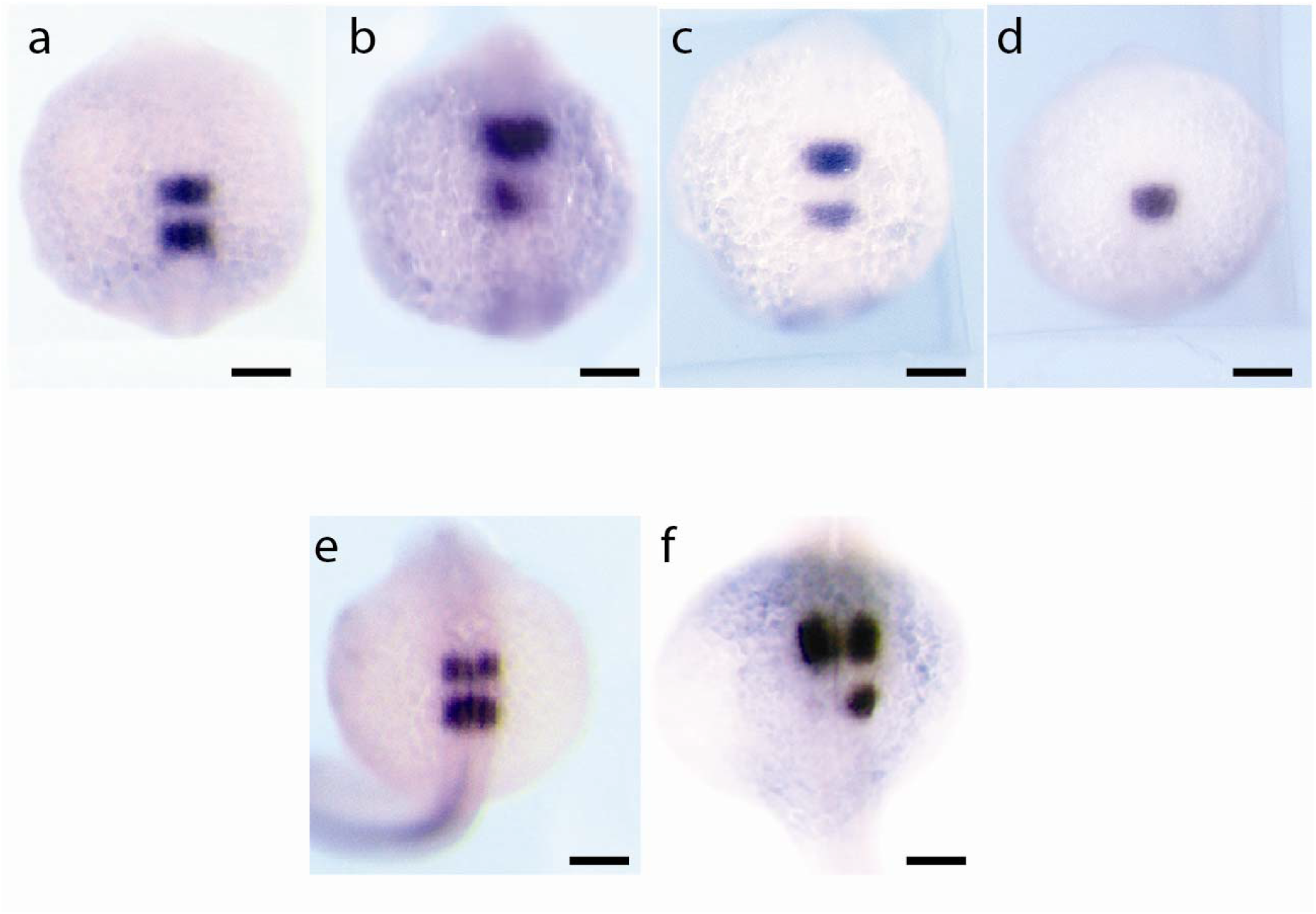
Inhibition of RA signaling disrupts bilateral symmetry of the hindbrain. (a-d) 8-10 somite zebrafish embryos hybridized *in situ* with a *krox20* probe labeling r3 and r5. a. 8-10s control embryo treated with 1% DMSO. b-d. 8-10s embryos treated with 10 μM BMS showing different types of abnormal *krox20* expression patterns. Scale bars: 100 μm (e-f) 18-19 somite zebrafish embryos hybridized *in situ* with a *krox20* probe labeling r3 and r5. (e) Control embryo treated with 1% DMSO. (f) Embryo treated with 10 μM BMS showing an asymmetric krox20 expression pattern. Scale bars: 100 μm Dorsal views, anterior to the top.

**Extended data Figure 2:**
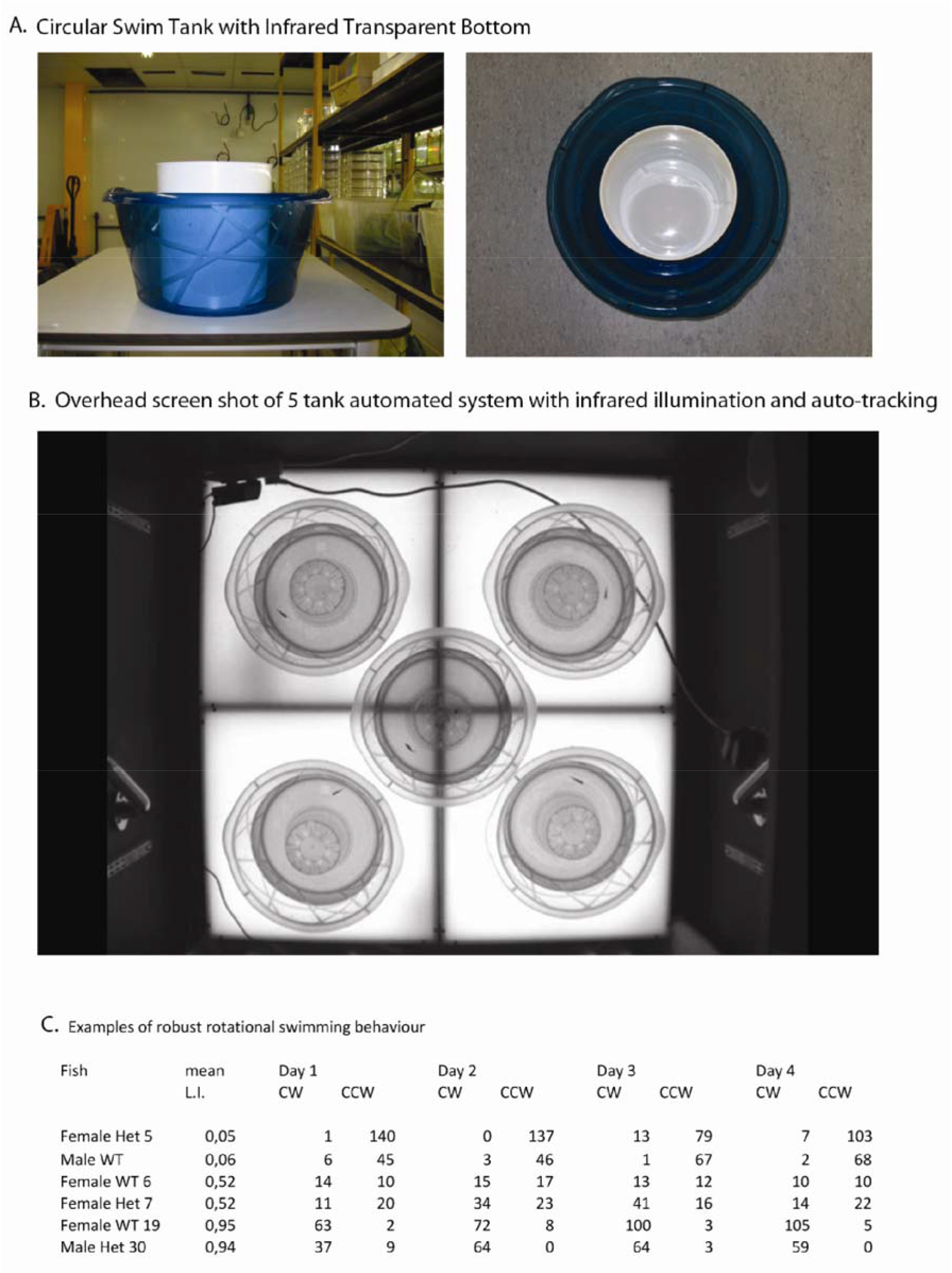
Behavioral setup for lateralization determination in adult zebrafish A. Example of the “donut”-shaped swimming chambers used during the test. B. Screen shot of the recording chambers, with the simultaneous infrared-based tracking of 5 fish. C. Examples showing individual adult fish with consistent directional swimming preferences from day to day over four consecutive days. Het =*bab^tb210^/+*; WT = wild-type +/+. Mean L.I. = Laterality Index averaged over all four days of the assay. CW = Clockwise; CCW = counterclockwise. Numbers under the column headings CW and CCW indicate the number of clockwise versus counter-clockwise circles that a given fish swam during a single 25 minute assay period on a single day.

**Extended data Figure 3:**
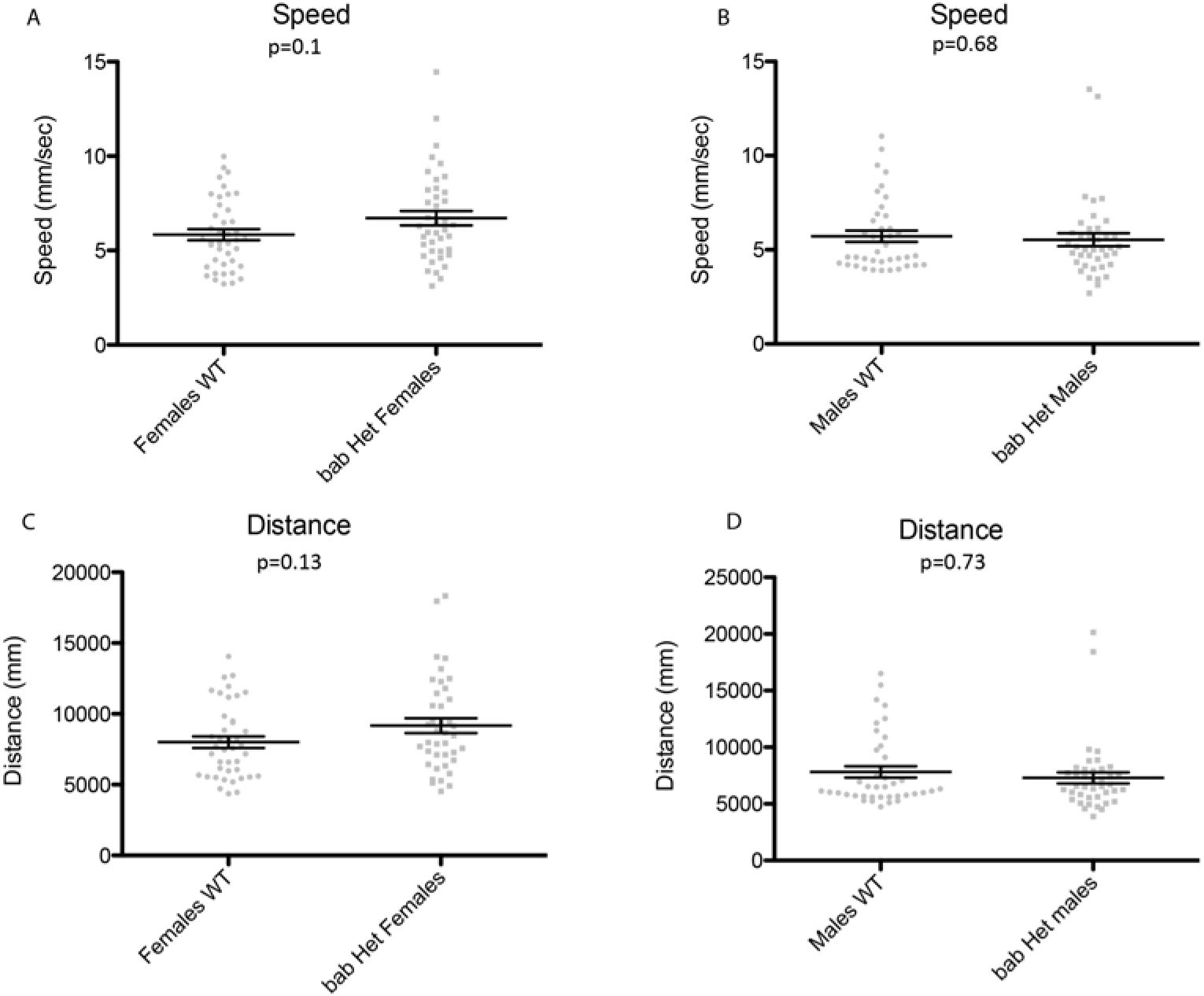
Swimming speed and swimming distance in *bab* heterozygotes (*bab^tb210^/+)* and wild type (+/+) adult zebrafish (a-b) Mean swimming speed during the rotation assays (25min recording), for the females (a, WT n= 41 and *bab* Het n= 40) and for the males (b, WT n=40 and *bab* Het n=40). (c-d) Mean total distance during the rotation assays for the females (c, WT n=41 and *bab* Het n=40) and for the males (d, WT n=40 and *bab* Het n=40). P values from Mann-Whitney test.

