## Supplementary Table 2 for "Rere-dependent Retinoic Acid signaling controls brain asymmetry and handedness"

Supplementary Table 2A. Asymmetric Krox r3 defects with MCC154

|  | n (at 8-10 s) | Sym Kr WT | Sym red r3 | Asym r3 | No r3 | Asym r5 | Sym DeltaC | Asym DeltaC |
| --- | --- | --- | --- | --- | --- | --- | --- | --- |
| Expt. 1 |  |  |  |  |  |  |  |  |
| 1% EtOH | 34 | 34 | 0 | 0 | 0 | 0 | 34 | 0 |
| 0.6 μM MCC154 | 47 | 3 | 40 | 4 | 0 | 0 | 47 | 0 |
| 6 μM | 40 | 0 | 29 | 9 | 2 | 0 | 36 | 4 |
| 64 μM | 44 | 0 | 29 | 13 | 2 | 0 | 44 | 0 |
| Expt.2 |  |  |  |  |  |  |  |  |
| 0. 4% EtOH | 11 | 11 | 0 | 0 | 0 | 0 | 9 | 2 |
| 0.25 μM | 33 | 33 | 0 | 0 | 0 | 0 | 28 | 5 |
| 2.5 μM | 43 | 1 | 34 | 8 | 0 | 0 | 41 | 2 |
| 25 μM | 31 | 1 | 15 | 15 | 0 | 0 | 26 | 5 |

PSM: Presomitic mesoderm. n: number of embryos. EtOH: ethanol. 8-10s: eight to ten somite stage. Sym Kr WT: wild-type *krox20* pattern in rhombomeres r(3) and r(5). Sym red r3: bilaterally symmetric but reduced *krox20* expression in r(3). Asym r3: Left-right asymmetric *krox20* expression in r(3). No r3: no obvious r(3) stripe of *krox20* expression. Sym and Asym *deltaC*: Bilaterally symmetric and left-right asymmetric *deltaC* expression in the PSM.

Supplementary Table 2B. Rare asymmetric rhombomere 3 defects in *giraffe ^rw716^*(*cyp26a1*) homozygotes.

|  | n total | Symmetric r3 | Sym reduced or lost r3 | Asymmetric r3 | Sym r5 |
| --- | --- | --- | --- | --- | --- |
| *Gir* inx 1 | 128 | 122 | 4 | **2** | 128 |
| *Gir* inx 2 | 104 | 101 | 1 | **2** | 104 |
| *Gir* inx 3 | 86 | 84 | 1 | **1** | 86 |

Krox 20 probe; 22-26 somites stage. All embryos showing an asymmetric Krox 20 r3 phenotype genotyped as *giraffe* homozygotes.
