## Supplementary Table 3 for "Rere-dependent Retinoic Acid signaling controls brain asymmetry and handedness"

**Supplementary Table 3: Nodal-dependent LR asymmetry is normal in *rerea* mutants and BMS-treated embryos**

| Crosses | Probe | Stage fixed | n | Phenotype |  |
| --- | --- | --- | --- | --- | --- |
| 4 pooled *bab ^tb210^*/+ incrosses | *southpaw* | 16-19s | 164 | % Left LPM  99 |  |
| WT AB cross† |  | 16-18s | 36 | 92 |  |
| WT AB cross† |  | 18-20s | 34 | 100 |  |
|  |  |  |  | % Left habenula* |  |
| 3 pooled *bab ^tb210^*/+ incrosses | *lefty1* | 21-22s | 83 | 99 |  |
| WT AB cross† |  | 19-22s | 28 | 100 |  |
| WT AB cross† |  | 21-23s | 24 | 100 |  |
| *bab ^tb210^*/+ incross | *uncx4.1* | 12-14s | 137 | % Symmetric Somites  99 |  |
| *bab ^tb210^*/+ incross  BMS 204493 Treatment | *uncx4.1* | 13-14s | 285 | 98 |  |
|  |  |  |  | % Left habenula* |  |
| DMSO 1% | *lefty1* | 24--25s | 11 | 82 |  |
| 10 μM BMS |  | 24--25s | 24 | 83 |  |
| 100 μM BMS |  | 24--25s | 17 | 94 |  |
| 0.1% DMSO | *lefty1* | 24-26hpf | 11 | 64 |  |
| 10 μM BMS204493 |  | 24-26hpf | 13 | 62 |  |
| BMS 204493 Positive Control |  |  |  |  |  |
|  |  |  |  | % WT (r3, r5) | % AbN |
| DMSO 1% | *krox 20* | 24--25s | 51 | 100 | 0 |
| 10 μM BMS |  | 24--25s | 41 | 49 | 51 |

†Table 3, Long et al. 2003.

n: number of embryos, s: somite. Hpf: hours post-fertilization.

% left LPM: % of embryos with *southpaw* expression in left but not right lateral plate mesoderm.

*% left habenula (left habenular nucleus of the epithalamus) was calculated from the formula: [(number of embryos with *lefty1(lft1)* expression only in the Left habenula) divided by (number of embryos with *lft1* only in the left habenula +number of embryos with *lft1* only in the right habenula + number of embryos with Bilateral (in both right and left habenula) *lft1* expression)] x 100

AbN: Abnormal *krox20* expression in hindbrain
