## Supplementary Table 4 for "Rere-dependent Retinoic Acid signaling controls brain asymmetry and handedness"

**Supplementary Table 4: Quantification of the number of embryos (homozygote *bab-/-* mutants, heterozygous *bab+/-* and wildtype siblings) showing symmetrical (sym) or asymmetrical (asym) patterns of r6 reticulospinal neurons (RS) or Mauthner cells (M), as revealed by retrograde labeling.**

L, R: missing reticulospinal neuron on the left or right side, respectively. *bab^tb210^ allele.*

| **Expt** | **bab/bab** | | | **bab/+** | | | **WT** | | |
| --- | --- | --- | --- | --- | --- | --- | --- | --- | --- |
|  | **No RS,**  **sym M** | **No RS,**  **asym M** | **total number** | **sym RS** | **asym RS** | **tot nb** | **sym RS** | **asym RS** | **total number** |
| Exp 1 | 8 | 0 | 8 | 7 | 5 (4L, 1R) | 12 | 5 | 1 (extra right RS) | 6 |
| Exp 2 | 11 | 1 | 12 | 14 | 11 (7L, 5R) | 25 | 6 | 1 (1L) | 7 |
